# Ribosomal protection as a linezolid resistance mechanism in *Mycobacterium abscessus*

**DOI:** 10.1101/2025.10.24.684387

**Authors:** Tobias Funck, Kerry McGowen, Mark R. Sullivan, Samuel Zinga, Ian D. Wolf, Dennis Nurjadi, Claudia M. Denkinger, Eric J. Rubin

## Abstract

*Mycobacterium abscessus* has emerged as a significant pulmonary pathogen characterized by its resistance to most first-line antimycobacterial drugs. Recent investigations have highlighted the clinical efficacy of including the oxazolidinone antibiotic linezolid in *M. abscessus* combination therapies, despite moderate resistance frequently being observed in patient isolates. Even with the potential usefulness of linezolid, the mechanisms that drive linezolid resistance in *M. abscessus* remain poorly understood. In several bacterial pathogens, including *Mycobacterium tuberculosis*, ATP-binding cassette (ABC) family proteins of the F subtype (ABC-F) have been found to confer antibiotic resistance to ribosome-targeting antibiotics, including linezolid. Here, we identified an *M. abscessus* ABC-F protein, MAB_2736c, that causes specific resistance to antibiotics that bind the 50S ribosomal subunit, including linezolid, macrolides, and chloramphenicol. These results demonstrate that targeting ABC-F proteins could help combat intrinsic resistance to several ribosome-targeting antibiotics in mycobacteria.

## Introduction

*Mycobacterium abscessus* is one of the few nontuberculous mycobacteria (NTM) species that can cause clinically significant infections in humans and is one of the pathogens responsible for the rise in opportunistic lung infections over the past two decades [1]. *M. abscessus* infection is notoriously difficult to treat, with cure rates estimated at less than 50% [2,3], primarily due to broad-spectrum antibiotic resistance [4]. Consequently, current treatment regimens for *M. abscessus* infections typically require combination therapies lasting 18–24 months [5]. This highlights the need for improved therapeutic options. The oxazolidinone antibiotic linezolid was recently shown to improve clinical outcomes when included in regimens to treat *M. abscessus* infections [6,7]. Additionally, linezolid substantially improved treatment success when included in multidrug regimens to treat tuberculosis, a disease caused by a closely related pathogen, *Mycobacterium tuberculosis* (Mtb). Linezolid is now included in the WHO guidelines for multidrug-resistant tuberculosis [8,9]. Since *M. abscessus* has limited therapeutic options and linezolid has conceivable potential to be included in regimens, it is imperative to better understand linezolid resistance in *M. abscessus*.

There are numerous routes that bacteria can take to achieve drug resistance. An increasingly recognized mechanism of resistance to ribosome-binding antibiotics involves F subtype ATP-binding cassette (ABC-F) proteins. The ABC superfamily is an ancient group of diverse proteins ubiquitous across all kingdoms of life. ATP-dependent transporters are the most well-known and well-characterized ABC family proteins, with many implicated in multidrug efflux [10,11]. However, several proteins involved in drug resistance containing an ABC cassette have recently been re-characterized from membrane transporters to ribosome-binding proteins. These ABC-F family proteins are located in the cytosol, where they bind directly to ribosomes and modulate their function, instead of transporting molecules across membranes [12,13]. Consequently, several ABC-F family proteins confer antibiotic resistance (ARE) by binding the ribosome to alleviate translational inhibition from antibiotics that target the large ribosomal subunit [12–17].

ARE-ABC-F proteins are categorized into three functional groups based on the antibiotics to which they confer resistance: (i) PLS_A_ for those that protect against pleuromutilins, lincosamides, and streptogramins A, (ii) MS_B_ for those that protect against macrolides and streptogramins B, and (iii) PhO for those that protect from phenicols and oxazolidinones [12–17]. All ARE-ABC-F proteins share two similar ABC nucleotide-binding domains that are connected by a helical linker called the antibiotic-resistance domain (ARD). The ARD has been shown to interact with the P-site tRNA in several ARE-ABC-F proteins. This allows the ARE-ABC-F protein to access the peptidyl transferase center, leading to a cascade of conformational changes that result in the displacement of antibiotics from their binding sites [14,18–20].

PLS_A_ and MS_B_ ARE-ABC-F proteins have been well-characterized across bacterial species, including *M. abscessus* [21]; however, PhO ARE-ABC-F proteins are less well-studied. Several PhO ARE-ABC-F proteins have recently been identified and characterized in other pathogenic species. For example, OptrA [22–25] and PoxtA [14,26,27], found in *Enterococcus* and *Staphylococcus* species, have been shown to be important antibiotic resistance factors with significant clinical implications. Furthermore, crystal structures of these proteins have revealed structural evidence for the ability of PhO ARE-ABC-F proteins to dislodge oxazolidinones and phenicols [14]. Additionally, recent work has demonstrated that OcrA (Rv1473) in Mtb is also an ARE-ABC-F PhO protein that confers resistance to linezolid [28]. These emerging pieces of evidence led us to investigate whether *M. abscessus* linezolid resistance is driven in part by the action of ABC-F family proteins.

## Results

### *MAB_2736c* likely encodes for an antibiotic resistance ABC-F

A homology search of amino acid sequences revealed that the *M. abscessus* gene *MAB_2736c* shares 84% sequence identity with that of *ocrA* (*Rv1473*) from Mtb [29,30]. To validate that *MAB_2736c* encodes an ABC-F family protein, its predicted tertiary structure was generated using AlphaFold [31], and this predicted structure was aligned with the predicted structure of OcrA as well as the X-ray crystal structure of PoxtA from *E. faecalis* [14] (Figures 1A–B). Structural alignment of MAB_2736c revealed a root mean square deviation (RMSD) of 1.709 Å relative to OcrA, supporting its classification as an ABC-F family protein. Although the RMSD relative to PoxtA is higher (4.581 Å), the two proteins share visually similar folds and conserved structural motifs (Figures 1A–B), further supporting a functional relationship despite the overall lower structural alignment.

**Figure 1.**
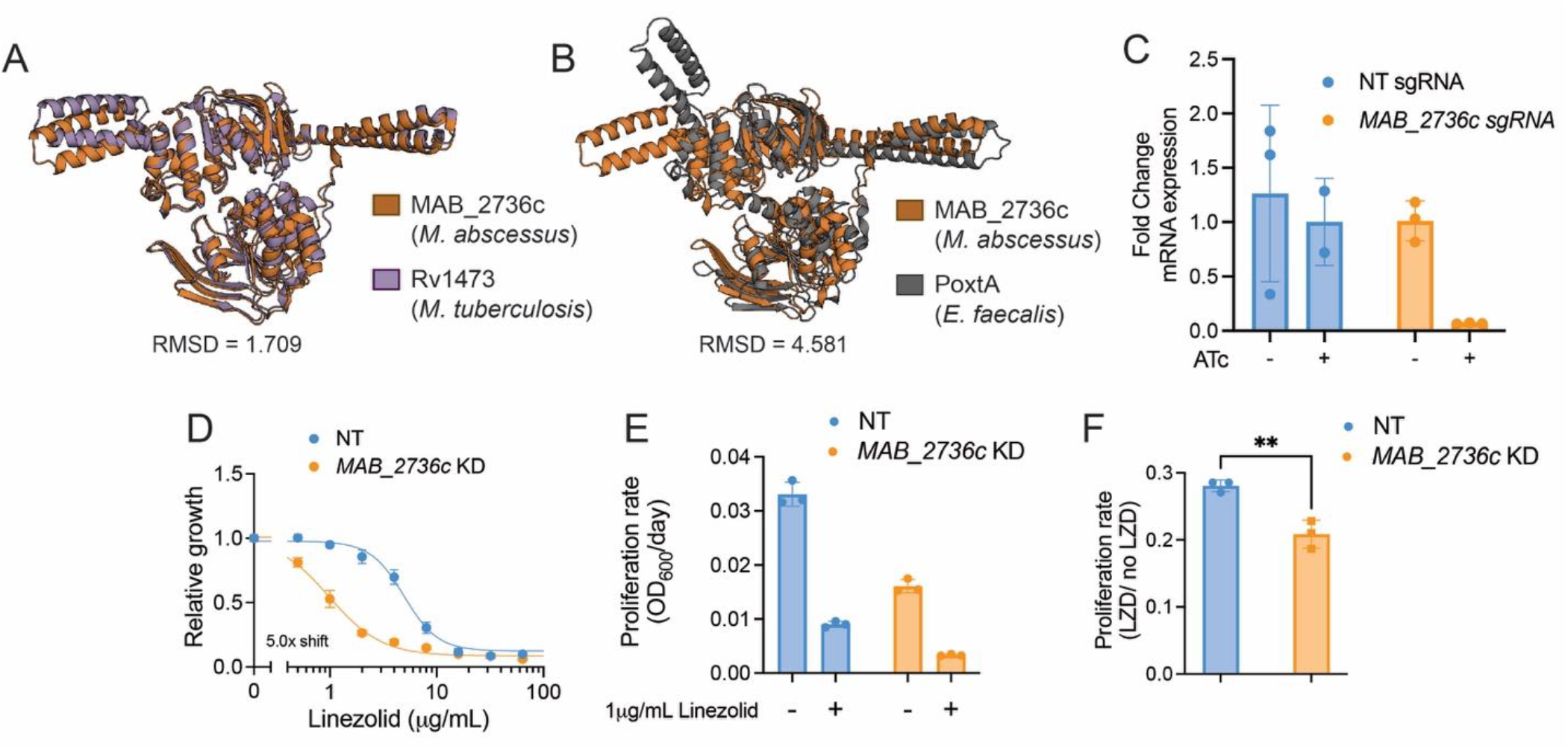
MAB_2736c knockdown displays increased sensitivity to linezolid. **(A)** Alignment of predicted protein structure of MAB_2736c from *M. abscessus* ATCC 19977 (orange) with predicted protein structure of Rv1473 from *M. tuberculosis* H37Rv (purple), and **(B)** PoxtA from *E. faecalis* crystal structure (grey). R.M.S.D = root mean square deviation. **(C)** Fold change mRNA expression of non-targeting (NT) sgRNA and *MAB_2736c* sgRNA in the presence or absence of 500 ng mL^-1^ ATc. **(D)** Relative proliferation as measured by resazurin reduction for *M. abscessus* ATCC19977 with CRISPRi construct targeting *MAB_2736c* or NT treated with indicated concentrations of linezolid in the presence of 500 ng mL^-1^ ATc. Values normalized to DMSO vehicle. **(E)** Proliferation rates calculated from OD600 measurements for a *MAB_2736c* knockdown (KD) and a non-targeting control (NT) in the presence of 500 ng mL^-1^ ATc in *M. abscessus* ATCC19977 grown in indicated concentration of linezolid for 48 hours. Values normalized to ATc-uninduced cells. **(F)** Ratio of proliferation rates under linezolid treatment normalized to vehicle treatment. p-value derived from unpaired, two-tailed t-test. For **(C-F)** *n* = 3 biological replicates, and data are presented as individual values along with mean ± s.d. NT = non-targeting. KD = knockdown. ATc = anhydrous tetracycline. LZD = linezolid.

We next sought to determine whether MAB_2736c might play a role in linezolid resistance. We utilized an anhydrotetracycline (ATc)-inducible Cas9/CRISPRi system in the *M. abscessus subspecies abscessus* reference strain (ATC19977) to determine whether depletion of this protein is sufficient to modulate linezolid susceptibility [32,33]. We found that the induced knockdown of *MAB_2736c* (Figure 1C) exhibits an increased sensitivity to linezolid (Figures 1D–F). Together, these data suggest that MAB_2736c is most likely an ARE-ABC-F protein and could play a role in antibiotic resistance similar to OcrA and other PhO proteins.

### Overexpression of *MAB_2736c* increases linezolid resistance

Though these results indicate that loss of MAB_2736c renders *M. abscessus* more susceptible to linezolid, those effects could be due to indirect translational defects downstream of MAB_2736c knockdown, as other ABC-F family proteins are known to modulate translation in a broad fashion, such as aiding in ribosome assembly [34]. To assess whether MAB_2736c can actively modulate linezolid efficacy, we next examined whether *MAB_2736c* overexpression confers increased linezolid resistance. Constitutive overexpression of *MAB_2736c* in the *M. abscessus* reference strain (Figure 2A) is sufficient to increase levels of resistance to linezolid (Figure 2B). While this result does not exclude the possibility of indirect translational effects, it is consistent with a direct role in linezolid protection. To determine whether this effect holds broadly true across mycobacteria, we also expressed *MAB_2736c* in *Mycobacterium smegmatis*, a nonpathogenic mycobacterial species. *M. smegmatis* also became significantly more resistant to linezolid with overexpression of *MAB_2736c* (Figures 2C–D), suggesting that *MAB_2736c* is sufficient to produce linezolid resistance through a mechanism that is generally applicable across mycobacteria.

**Figure 2.**
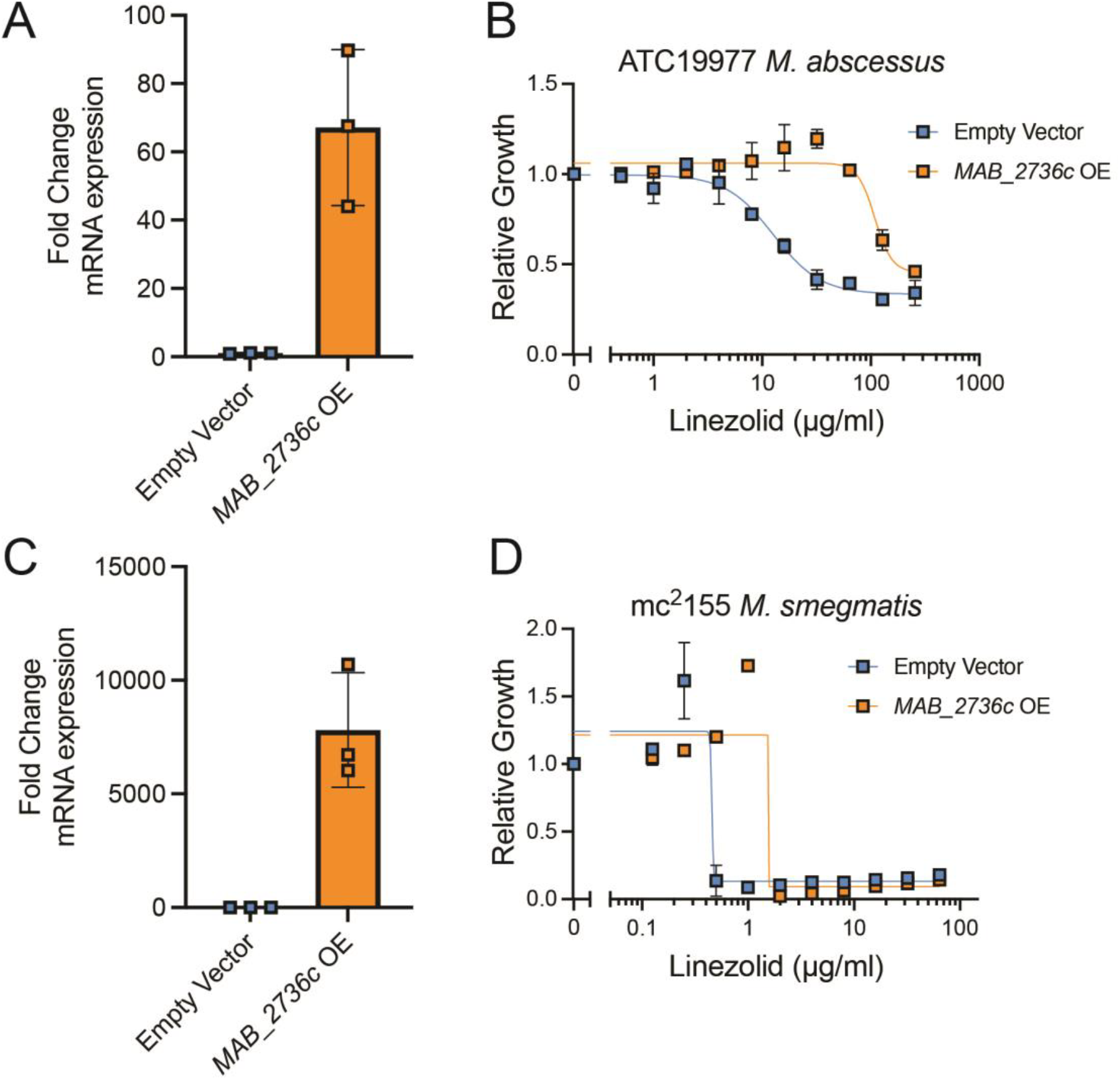
MAB_2736c overexpression is sufficient to cause linezolid resistance. Fold change mRNA expression of empty vector control and *MAB_2736c* overexpressing (OE) vector in **(A)** ATC19977 *M. abscessus* and **(C)** mc^2^155 *M. smegmatis*. Relative proliferation as measured by resazurin reduction for parental strains with an empty vector or strains overexpressing MAB_2736c vector (*MAB_2736c* OE) of **(B)** *M. abscessus* ATCC19977, and **(D)** *M. smegmatis* mc^2^155 exposed to the indicated concentrations of linezolid. Data are presented as individual values along with mean +/− SD. n=3 biological replicates.

### MAB_2736c changes the susceptibility to multiple ribosome-targeting antibiotics

All previously characterized ARE-ABC-F proteins provide resistance to a certain subset of antibiotics that target the 50S subunit of the ribosome [12–17]. Consistent with this observation, *MAB_2736c* knockdown induces increased sensitivity to chloramphenicol (Figure 3A), a compound whose effect is also modulated by OcrA [28] and other PhO ARE-ABC-F proteins [14,22–27]. Unexpectedly, *MAB_2736c* knockdown also induces sensitivity to the macrolides erythromycin and clarithromycin (Figures 3B– C). This was a notable finding given that other PhO ARE-ABC-F proteins, like PoxtA and OptrA, do not alter sensitivity to macrolides [14]. However, earlier work, when OcrA was thought to be an ABC transmembrane drug exporter, showed that *ocrA* (*rv1473*) mutants in Mtb also have an increased susceptibility to macrolides [35]. These observations suggest that MAB_2736c may influence resistance more broadly than previously characterized PhO ARE-ABC-F proteins or that MAB_2736c is promoting drug resistance through a different mechanism.

**Figure 3.**
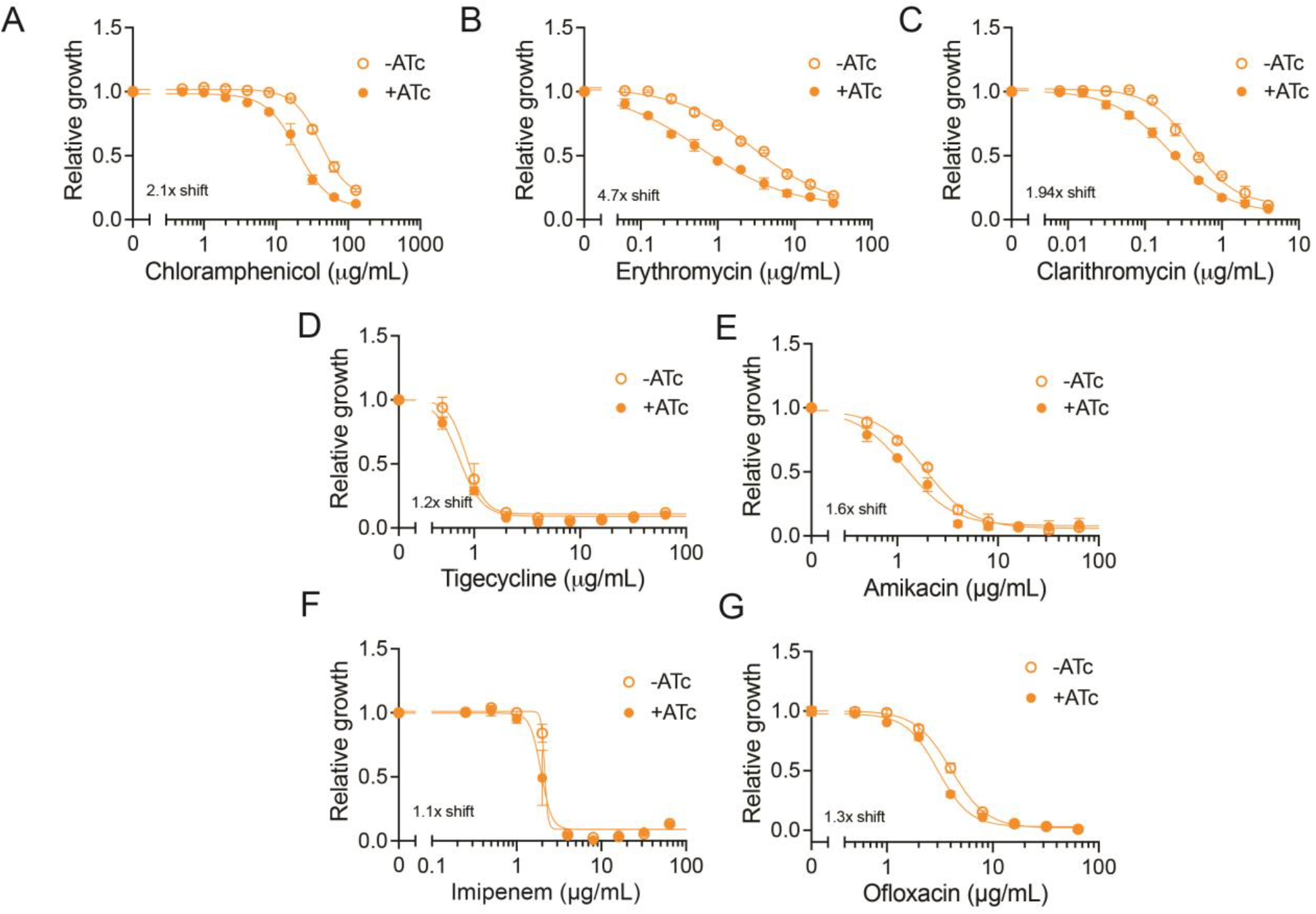
MAB_2736c knockdown mutant is sensitive to multiple 50S ribosome-targeting antibiotics. Relative proliferation as measured by resazurin reduction for *M. abscessus* ATCC19977 with CRISPRi construct targeting *MAB_2736c* treated with indicated concentrations of **(A)** chloramphenicol, **(B)** erythromycin, **(C)** clarithromycin, **(D)** tigecycline, **(E)** amikacin, **(F)** imipenem, and **(G)** ofloxacin in the presence or absence of 500 ng mL^-1^ ATc. Values normalized to DMSO vehicle. n= 3 biological replicates. Data are presented as individual values along with mean ± s.d. ATc = anhydrous tetracycline.

To distinguish these possibilities, we examined whether *MAB_2736c* knockdown affects sensitivity to antibiotics that target translation through the 30S subunit (amikacin and tigecycline) as well as antibiotics that target DNA (ofloxacin) or cell wall synthesis (imipenem), which are not expected to be affected by the action of traditional PhO ARE ABC-F proteins. *MAB_2736c* knockdown does not alter sensitivity to these antibiotics (Figures 3D–G), suggesting that MAB_2736c provides specific protection against translation inhibitors that target the 50S subunit, similar to other PhO ARE-ABC-F proteins. However, the observed broader range of protection, including macrolides, raises the possibility that MAB_2736c operates through a distinct or partially divergent mechanism compared to other PhO ARE-ABC-F proteins.

### MAB_2736c protects purified ribosomes against linezolid

To further characterize the mechanistic effects of MAB_2736c expression on ribosome function, we sought to evaluate the effects of MAB_2736c on translation in a cell-free transcription–translation system that utilizes ribosomes derived from *Escherichia coli* [36]. We simultaneously expressed a fluorescent reporter protein, mNeonGreen, with either MAB_2736c or a control protein from *E. coli*, dihydrofolate reductase (DHFR) (Figure 4A). Following 30 minutes of undisturbed translation of MAB_2736c or DHFR, we then added linezolid and monitored whether MAB_2736c protected translating ribosomes (Figure 4A). In the absence of drug, MAB_2736c reduces the fluorescent reporter translation baseline rate compared to either DHFR or the no template control (Figures 4B–C), suggesting that MAB_2736c may modulate or interfere with *E. coli* ribosome function *in vitro*. The addition of linezolid decreases the rate of translation as expected in the control template condition but only produces a modest decrease in translation rate in the MAB_2736c expressing condition (Figures 4B–C). To account for the different baseline rates of translation, we normalized the protein synthesis rate to vehicle-treated (DMSO) and found that MAB_2736c displays modest protection of translation compared to DHFR when treated with 25 μM linezolid (Figure 4D). This protective effect is diminished at higher doses of linezolid (Figure 4D), consistent with the observation that more saturating doses of linezolid are able to inhibit *M. abscessus* growth. These results demonstrate that MAB_2736c is likely able to provide some direct ribosomal protection against linezolid and suggest a role for ribosome protection in *M. abscessus* linezolid resistance.

**Figure 4.**
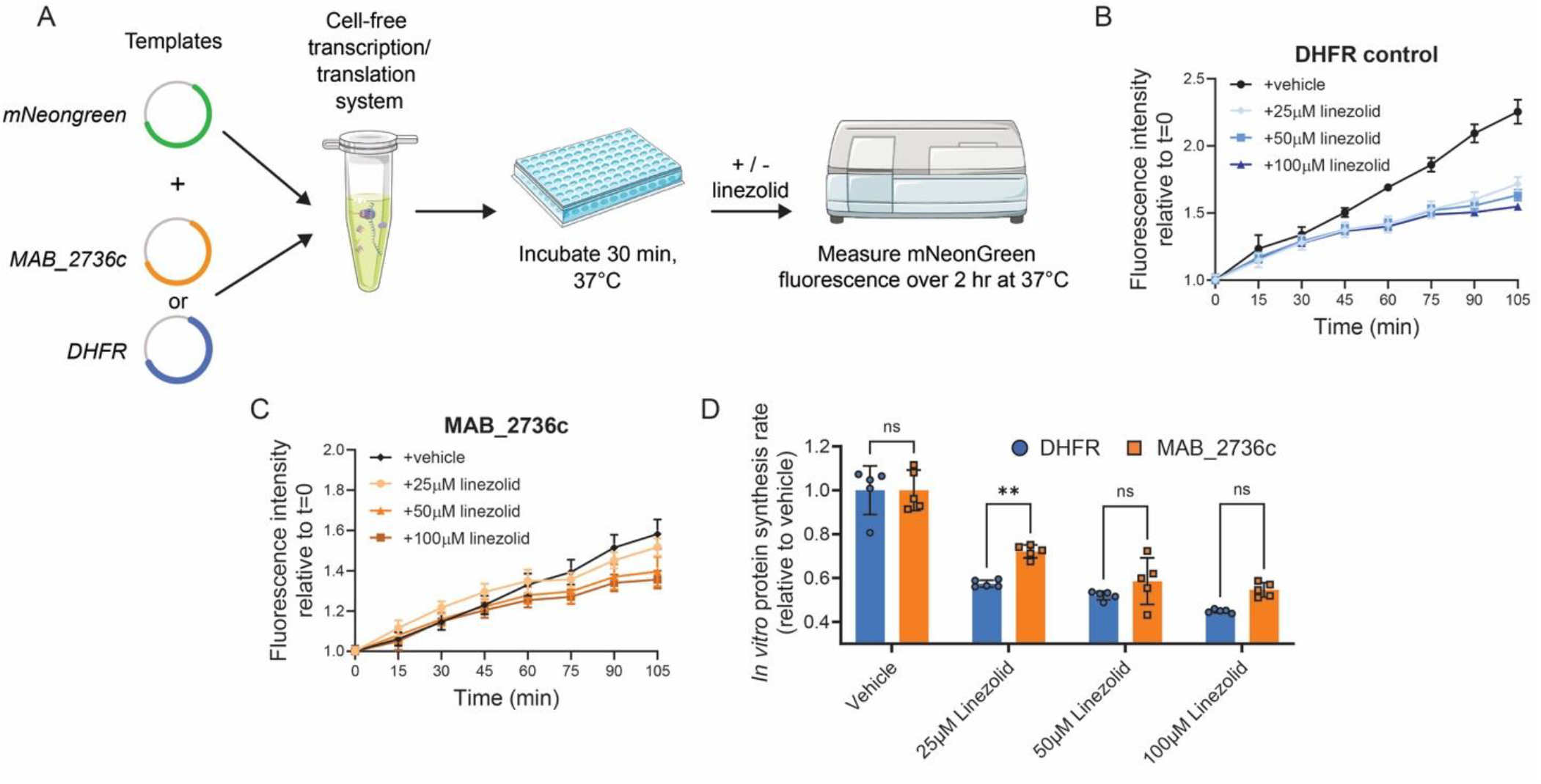
MAB_2736c protects purified ribosomes against linezolid. **(A)** Schematic of cell-free transcription/translation assay derived from *E. coli*. DHFR = Dihydrofolate reductase. **(B)** mNeonGreen fluorescence when co-expressed with DHFR in the presence 25μM, 50μM, 100 μM linezolid or DMSO. n=5 independent samples. **(C)** mNeonGreen fluorescence when co-expressed with MAB_2736c in the presence 25μM, 50μM, 100 μM linezolid or DMSO. n=5 independent samples. **(D)** Relative protein synthesis rate for mNeonGreen co-expressed with MAB_2736c or DHFR in the presence of 25μM, 50μM, or 100μM linezolid, or DMSO. Rates were calculated from post-linezolid addition until 105 minutes elapsed. Data are presented as individual values along with mean +/− SD. p-values derived from Šídák’s multiple comparisons test after two-way ANOVA. n=5 biological replicates.

## Discussion

Numerous ABC-F proteins have been shown to confer antibiotic resistance in several pathogenic species [12–17]. However, their roles in pathogenic mycobacteria remain largely unexplored. In this study, we identified MAB_2736c in *M. abscessus* as an ARE-ABC-F protein that confers resistance to linezolid, chloramphenicol, and macrolides, likely through a ribosome protection mechanism.

Typically, ARE-ABC-F proteins are categorized into functional groupings based on antibiotics to which they confer resistance. For example, ARE-ABC-F proteins that confer resistance to linezolid and chloramphenicol belong to the PhO group of ARE-ABC-F proteins [14]. However, in contrast to other PhO ARE-ABC-F proteins, MAB_2736c expression also alters sensitivity to other 50S ribosomal subunit-targeting antibiotics, including macrolides. Our findings suggest that MAB_2736c could act in a mechanistically distinct manner from other PhO proteins. This distinction highlights the need to re-evaluate the current classification system for ARE-ABC-F proteins. Furthermore, we find a bacterial growth deficit caused by MAB_2736c depletion even in the absence of antibiotic pressure, hinting at an integral biological role of MAB_2736c beyond drug resistance. Further investigation is required to discern how MAB_2736c modulates translation under baseline conditions. Structural insights into how MAB_2736c binds to the mycobacterial ribosome could help illuminate how MAB_2736c differs from other ARE-ABC-F family proteins and provide insight into its native functions. Consequently, this could also help inform targeted drug design for MAB_2736c.

Inhibition of MAB_2736c may enhance the efficacy of multiple clinically relevant ribosome-targeting antibiotics. Further, several avenues to overcome the antibiotic resistance caused by ARE-ABC-F proteins have been proposed; these include the development of ribosome-targeting drugs with higher ribosomal affinity to outcompete ABC-F proteins, as well as ABC-F protein binding site mimics to inhibit ribosome function to stall further translation [17]. Given the broad effects of MAB_2736c on the potency of several classes of ribosome-targeting antibiotics, applying these approaches in *M. abscessus* might be a valuable way to improve our ability to target bacterial translation and combat resistance. Since our findings were obtained using genetically manipulated laboratory strains, future studies in clinical isolates or in vivo models will be needed to validate the clinical relevance of targeting MAB_2736c and to determine whether they translate to improved treatment outcomes. Overall, this work advances our understanding of ARE-ABC-F proteins in mycobacteria and supports previous notions that ARE-ABC-F proteins may represent attractive drug targets to potentiate the effects of current antibiotics.

## Acknowledgements

T.F. was supported by a Boehringer Ingelheim Fonds MD fellowship. M.R.S. received support as a Merck Fellow of the Damon Runyon Cancer Research Foundation, DRG-2415-20. S.Z. was supported by award numbers T32GM145407, T32GM007753 and T32GM144273 from NIGMS and award number F30AI188797 from NIAID. E.J.R. was supported by NIH/NIAID under award number R01AI179642. The content is solely the responsibility of the authors and does not necessarily represent the official views of the National Institute of General Medical Sciences, the National Institute of Allergy and Infectious Diseases, or the National Institutes of Health.

## Author contributions

Conceptualization: K.M., T.F., M.R.S.; Methodology development: K.M., T.F., M.R.S.; Experimentation: K.M., T.F., S.Z., I.D.W., M.R.S.; Formal analysis: K.M., T.F., M.R.S.; Visualization: K.M. T.F.; Original draft: K.M., T.F.; Second draft review and edits: K.M., T.F., M.R.S.; Funding acquisition: T.F., M.R.S., E.J.R.; Supervision: D.N., C.M.D., E.J.R., M.R.S.

## Data availability

All data supporting the findings of this study are available within the article.

## Methods

### Strains

All experiments were performed in the *Mycobacterium abscessus* subspecies *abscessus* type strain (ATCC19977) unless otherwise indicated. The experiments with *Mycobacterium smegmatis* utilized the strain mc^2^155 (ATCC700084). All plasmid construction was performed in *Escherichia coli* DH5α.

### Oligonucleotides

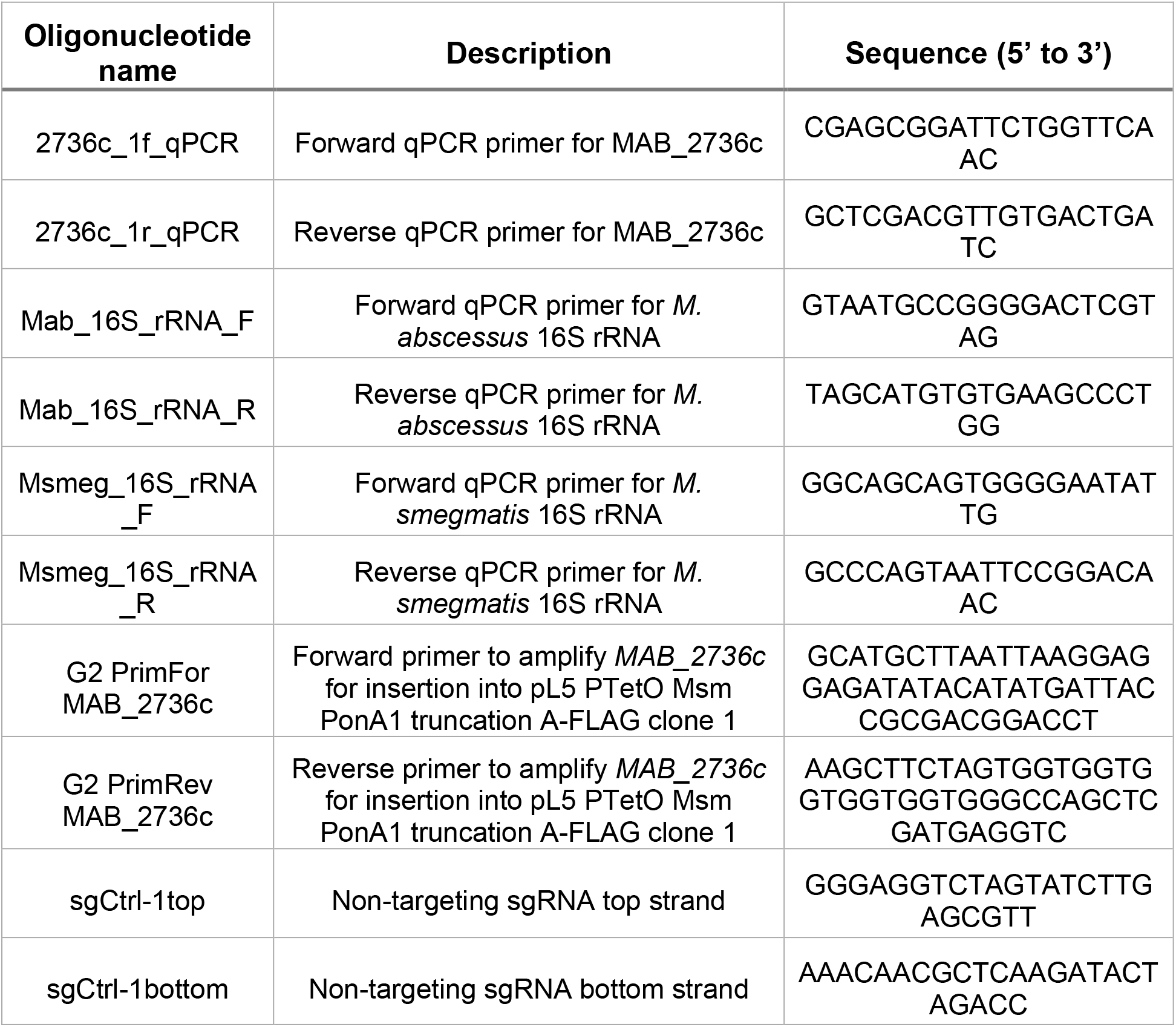
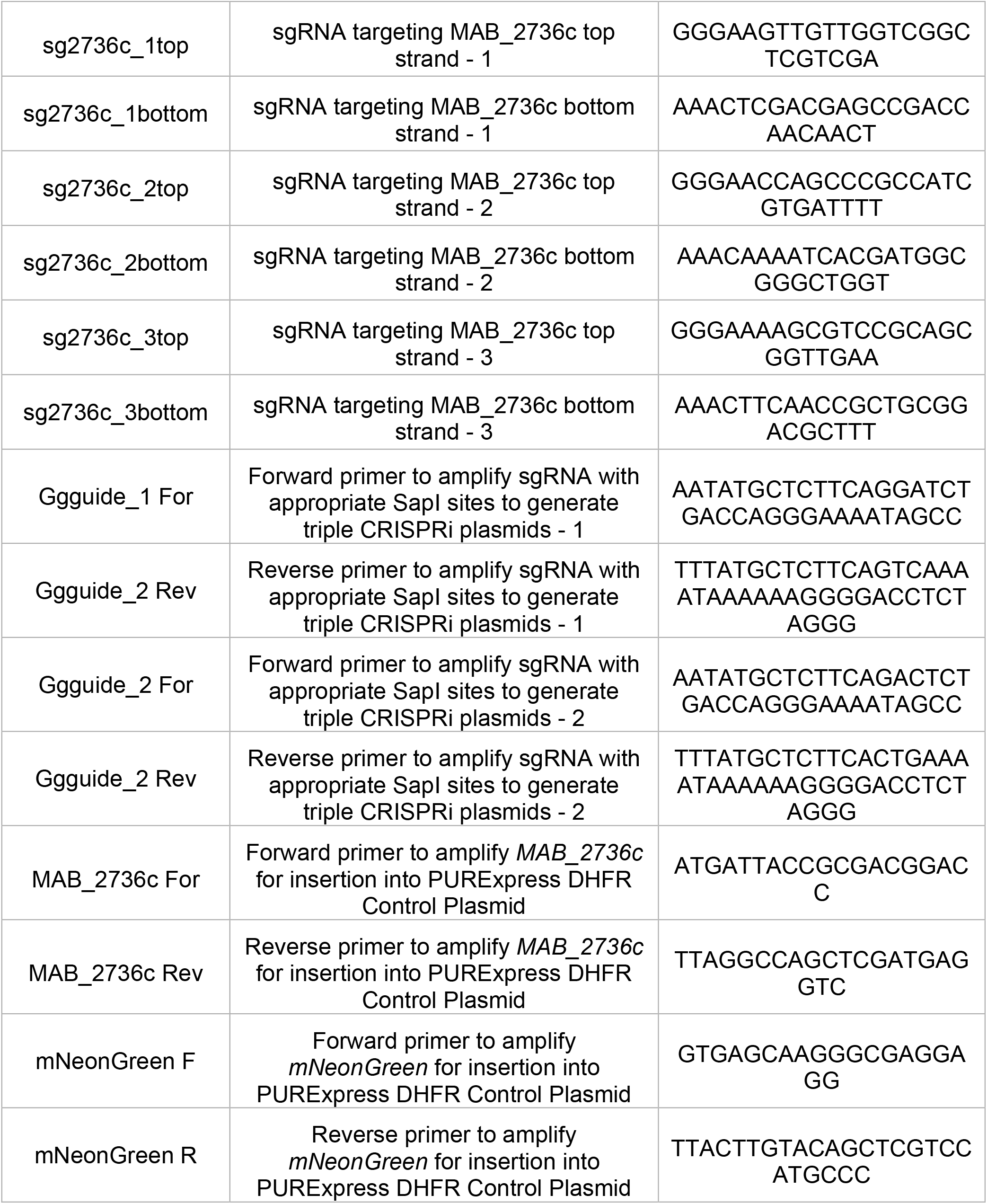

### Mycobacterial culturing conditions

All *M. abscessus* and *M. smegmatis* cultures were grown at 37°C in Middlebrook 7H9 broth (271310, BD Diagnostics) with 0.2% (v/v) glycerol (GX0185, Supelco), 0.05% (v/v) Tween-80 (P1754, MilliporeSigma), and 10% (v/v) oleic acid-albumin-dextrose-catalase (OADC) (90000-614, VWR) (*M. abscessus*) or albumin dextrose catalase (ADC) (*M. smegmatis*) with appropriate antibiotics.

### Mycobacterial transformations

*M. abscessus* or *M. smegmatis* were grown to an optical density (OD_600_) of 0.8, then washed thrice with sterile 10% glycerol by pelleting at 5000 x *g* for 7 minutes at 22°C. After final wash, cells were resuspended in 1% of the initial culture volume in 10% glycerol. 50 μL of electrocompetent mycobacteria were mixed well with 100 ng DNA in 2 μL water and then transferred to a 2 mm electroporation cuvette (89047-208, VWR). The cells were electroporated at 2500 V, 125 Ω, 25 μF using an ECM 630 electroporator (45-0651, BTX). 1 mL 7H9 + OADC broth was added to the electroporated cells, and cells recovered for 4 hours shaking at 150 rpm at 37°C. 100 μL of recovered cells were spread on 7H10 + 0.5% (v/v) glycerol + 10% (v/v) OADC agar plates with 50 μg ml^−1^ kanamycin sulfate using 4 mm borosilicate glass beads. Plates were incubated at 37°C for 4 days. For *M. smegmatis* strains a similar method was used, but with a shortened recovery time of 2h after electroporation and plates were incubated for only 3 days.

### Generation of mutant MAB_2736c strains

CRISPRi MAB_2736c plasmid was constructed using Addgene plasmid 166886 as previously described [33]. Briefly, plasmid 166886 was digested with BsmBI-v2 (NEB R0739L) and then gel purified. Three sgRNAs were designed to target three different locations of the non-template strand of MAB_2736c. Each individual sgRNA with appropriate overhangs was annealed and ligated (T4 ligase NEB M0202M) into three separate BsmBI-v2 digested backbones. To generate a plasmid with all three sgRNAs, a SapI-based Golden Gate cloning site 3′ to the first sgRNA scaffold was used as previously described [33]. Successful plasmid construction was verified using long-read sequencing (Plasmidsaurus). Non-targeting (NT) control was constructed in a similar manner but with scrambled, non-targeting sgRNAs. Triple MAB_2736c or NT plasmids were transformed into ATCC19977 and selected on 7H10 + 0.5% (v/v) glycerol + 10% (v/v) OADC agar plates containing 50 μg ml^−1^ kanamycin sulfate.

MAB_2736c constitutive plasmid was constructed into pL5 PTetO Msm PonA1 truncation A-FLAG clone 1 [37] using NdeI (R0111, NEB) and HindIII-HF (R3104, NEB) restriction digest followed by isothermal assembly with Phusion High-Fidelity Polymerase (M0530, NEB) and transformed into ATCC19977 and *M. smegmatis*. Successful transformants were selected on 7H10 + 0.5% (v/v) glycerol + 10% (v/v) OADC agar plates containing 50 μg ml^−1^ kanamycin sulfate.

### Minimum inhibitory concentration determination

*M. abscessus* and *M. smegmatis* were grown until mid-log phase (OD_600_ of 0.6-0.8). For CRISPRi experiments, cultures were induced for knockdown 18-24 hours prior to start of the assay with 500 ng μl^-1^ ATc in DMSO. Cultures were then diluted to OD_600_ of 0.003 and 200 μl aliquots were plated in technical triplicate in wells (3370, Corning) containing specified antibiotics or 500 ng μl^-1^ ATc when relevant. Antibiotic stocks were made as follows: 20 mg mL^-1^ linezolid (PZ0014, Sigma-Aldrich) in DMSO, 10 mg mL^-1^ clarithromycin (C9742, Sigma-Aldrich) in DMSO, 10 mg mL^-1^ amikacin disulfate salt (A1774, Sigma-Aldrich) in water, 10 mg mL^-1^ ofloxacin (O8757, Sigma-Aldrich), 1 mg mL^-1^ imipenem monohydrate (I0160, Sigma-Aldrich) in water, 20 mg mL^-1^ chloramphenicol (C0378, Sigma Aldrich) in ethanol, or 10 mg mL^-1^ tigecycline hydrate (PZ0021, Sigma Aldrich) in DMSO. The cells were then incubated at 37°C with shaking at 150 r.p.m. for 24 hours. 0.002% resazurin (R7017, Sigma Aldrich) in ddH2O was then added to each well and plates were incubated for an additional 24 hours at 37°C with shaking at 150 r.p.m. MIC determination was conducted using a Tecan Spark 10M plate reader (Mannedorf, Switzerland) by measuring absorbance at 570 nm and 600 nm and normalizing the ratio to background and the no drug control.

### Growth curve

CRISPRi strains were grown until mid-log phase (OD_600_ of 0.6-0.8) and pre-depleted with ATc at 500 ng mL^-1^ for 18-24 hours. Cultures were then back-diluted at a final OD_600_ of 0.02 and 200 μl of diluted cells were added in technical triplicate with DMSO or linezolid at 0.5 μg mL^-^1 and fresh ATc at 500 ng mL^-1^. Growth was determined by continuous OD_600_ measurement in 15-min intervals in a Spark 10M plate reader for 48 hours at 37°C with continuous shaking at 1000 rpm. Growth curve data were analyzed using Microsoft Excel 365 and GraphPad Prism 9.

### Protein structure alignments

Tertiary protein structures for MAB_2736c from *M. abscessus* ATCC19977 and OcrA from *M. tuberculosis* H37Rv [29,30] were predicted using AlphaFold [31]. MAB_2736c predicted protein structure was aligned with the predicted structure of OcrA and the X-ray crystal structure of PoxtA from *E. faecalis* [14] using PyMOL 3.0 [38]. RMSD scores (root mean square deviation) were determined using the alignment function within PyMOL 3.0.

### Quantitative PCR

For the *MAB_2736c* CRISPRi strain, cultures were grown in biological triplicate to mid-log phase, diluted back in +/-500 ng/ml ATc, and grown for 18-24 hr to achieve target knockdown. For the *MAB_2736c* overexpression strains in *M. abscessus* and *M. smegmatis*, cultures were grown in biological triplicate under kanamycin selection to an OD_600_ of 0.6-0.8. 2 OD_600_ equivalents of cells from each culture were harvested by centrifugation, resuspended in TRIzol (Thermo Fisher), and lysed by bead beating (Lysing Matrix B, MP Biomedicals) 4 x 4000 rpm. Total RNA was extracted from the TRIzol aqueous phase using column-based purification (R1018, Zymo Research). Residual genomic DNA was digested by addition of TURBO DNase buffer (AM2238, Thermo Fisher Scientific) (final concentration of 10%) and TURBO DNase (final concentration of 2%) for 1 hr at 37°C, then samples were purified with RNA clean-up columns (R1018, Zymo Research). cDNA was prepared using random hexamers (Life Technologies Superscript IV), column purified (28106, Qiagen) and then quantified by RT-qPCR on a QuantStudio 7 Flex real-time PCR machine (Thermo Fisher Scientific) using iTaq Universal SYBR Green Supermix (BioRad). All qPCR primer pairs were confirmed to display a linear response to cDNA concentration over a 64-fold range covering the experimental values. Relative cDNA abundance was calculated using the ΔΔCt method through normalization to the –ATc condition and to expression of the 16S rRNA.

### PURExpress *in vitro* protein synthesis

Plasmids carrying MAB_2736c or mNeonGreen compatible with the PURExpress in vitro protein synthesis kit were cloned into the manufacturer’s PURExpress DHFR Control Plasmid (E6800L, NEB) by swapping out DHFR with MAB_2736c (PCR amplified from ATC19977 gDNA) or mNeonGreen using standard restriction cloning with NdeI (R0111S, NEB) and PacI (R0527S, NEB). PURExpress reactions were set up following the manufacturer’s protocol with either no template, 200 ng pDHFR or 200 ng pMAB_2736c as well as 200 ng of mNeonGreen included in every reaction. Mixed reactions were aliquoted into black bottom 384–well plates (CLS3573, Corning), and the plates were incubated at 37°C for 30 minutes. 100 μM, 50 μM, 25 μM linezolid or DMSO control were spiked into appropriate wells. Plates were incubated for an additional 2 hours at 37°C in Spark 10M plate reader with ex/em of 506 nm/517 nm read at 15-minute intervals. PURExpress data were analyzed using Microsoft Excel 365 and GraphPad Prism 9.

### Statistical methods

For all data points, error bars represent the standard deviation of the y-variable on the graph. Statistical significance between two independent groups was queried with unpaired, two-tailed t-tests. For analyses involving multiple groups and experimental factors, statistical significance was assessed using two-way ANOVA followed by Šídák’s multiple comparisons test to adjust for multiple hypothesis testing. All statistical tests used are indicated in figure legends.

### Visualization

Schematic visualizations of experiments were created with Inkscape (version 1.4.2; Inkscape Project, https://inkscape.org). Illustrative elements were adapted from Servier Medical Art, which is licensed under a Creative Commons Attribution 4.0 International License (CC BY 4.0) (https://smart.servier.com). Statistical figures were generated with GraphPad Prism 9.

